# Zero-shot neural decoding of visual categories without prior exemplars

**DOI:** 10.1101/700344

**Authors:** Thomas P. O’Connell, Marvin M. Chun, Gabriel Kreiman

## Abstract

Decoding information from neural responses in visual cortex demonstrates interpolation across repetitions or exemplars. Is it possible to decode novel categories from neural activity without any prior training on activity from those categories? We built zero-shot neural decoders by mapping responses from macaque inferior temporal cortex onto a deep neural network. The resulting models correctly interpreted responses to novel categories, even extrapolating from a single category.

---

Neural decoding approaches typically train machine learning classifiers on responses to a set of stimuli and subsequently test the classifier using either different repetitions of the same training stimuli or responses to different exemplars from the same training categories. These approaches have been extremely successful in a wide variety of domains [1], but show limited generalization.

Zero-shot neural decoding, or interpreting neural activity without prior exposure to any similar information [2–6], holds great promise to improve the generalizability of neural information processing models. While standard decoders predict information directly from patterns of neural activity, zero-shot decoders map neural activity to an intermediate representation that constitutes a computational hypothesis for the neural code [2]. The intermediate representation is selected such that it has a known or easily learned relationship to a wide variety of to-be-predicted outputs. In an impressive recent demonstration of zero-shot decoding, Anumanchipalli and colleagues [6] reconstructed recognizable human speech from electrophysiological recordings in human motor cortex via a computational model of articulatory movement. Even though the decoding model was only trained to map neural activity to the articulatory model, and not representations of words or semantics, the models could reconstruct intelligible human speech. Here, we demonstrate such zero-shot decoding from electrophysiological responses for visual objects.

Beyond a feat of engineering, the degree of generalization has important consequences for the conclusions that can be drawn from a model of neural information processing. The greater the generalization, the stronger the evidence that a model captures generic processing beyond any particular set or class of stimuli. As an example, consider a standard linear decoder trained to distinguish whether responses along the ventral stream were evoked by images of airplanes or chairs. The decoder could interpolate within its training space to label neural responses to new images of airplanes or chairs, but it would not be able to accurately label neural responses to cars or tables. A zero-shot model can capture generic visual information and extrapolate to new categories on which it was not trained.

Constructing generic zero-shot decoders for visual objects necessitates a model for visual processing in the primate brain. How well do we understand the neural code for visual object processing? Deep convolutional neural networks (DCNNs) constitute a promising initial approximation to the cascade of computations along the ventral stream that support visual object recognition [7–11]. DCNNs are goal-directed, hierarchical, image-computable models capable of recognizing complex, natural objects and scenes [12], and representations in DCNNs predict object-evoked neural activity in *rhesus macaque* inferior temporal cortex (IT) [13–15], which is at the top of the ventral visual stream hierarchy and plays a central role in visual object recognition [16, 17].

While DCNNs are powerful pattern extractors, it remains possible that their performance predicting IT responses is driven by generalization within stimuli (e.g., different views of the same chair) or within categories (e.g., one type of chair to another). To test whether DCNNs capture the type of flexible visual processing accomplished by biological vision, the mapping from DCNNs to IT should generalize across object categories (e.g. chairs to cars). While some studies have shown extrapolation across categories [3, 13], the degree of generalization remains unclear. In an extreme case, can IT to DCNN mappings learned from neural activity evoked by a single object category extrapolate to novel categories? If the mappings generalize to new images from the same category, we can conclude that DCNN responses capture category-level information within IT. If the mappings generalize to new images from novel categories, this suggests that DCNNs capture generic visual information in IT beyond any one category.

To determine whether representations in DCNNs capture generic visual processing in the primate brain, we built zero-shot neural decoders for object category from multi-electrode array recordings in *rhesus macaque* IT (**Fig. 1a**). IT responses were evoked by images of computer-generated objects on natural scene backgrounds with high variation in position, size, and orientation (**Fig. 1b**). We tested whether zero-shot decoders trained on neural responses to a set of categories (e.g., airplane and chair images) can accurately label neural activity evoked by novel categories (e.g., cars and tables). In the most extreme instance, we tested whether zero-shot decoders trained on neural responses from a single category can generalize to label neural responses evoked by seven novel categories.

**Figure 1.**
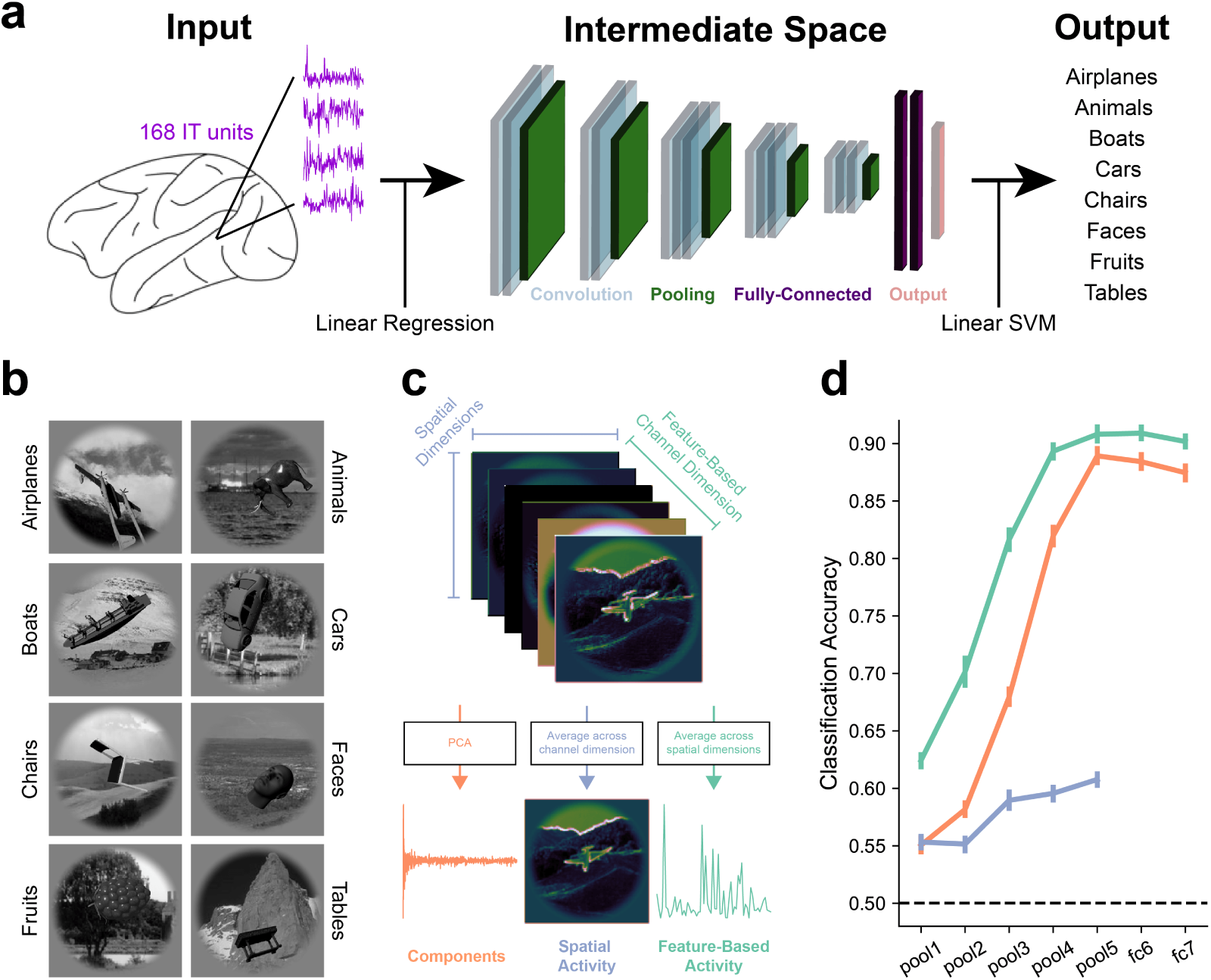
Overview of zero-shot approach and deep convolutional neural network (DCNN) architecture. **a**. Overview of zero-shot decoding pipeline. IT recordings were mapped to an intermediate space defined as unit activity in a deep convolutional neural network trained for object categorization. Pre-learned mappings from DCNN unit activity to object categories were used to generate predictions from DCNN-aligned IT recordings. The decoders are zero-shot if neural recordings from the test categories are withheld when learning the IT to DCNN. **b**. Example images from the eight object categories: Airplanes, Animals, Boats, Cars, Chairs, Faces, Fruits, Tables [18]. **c**. VGG-16 trained for object categorization on the ImageNet dataset [19] was used as the intermediate basis-space to which IT recordings were aligned. In the pooling layers, DCNN activity is organized along two spatial dimensions and a feature-based channel dimension. To summarize the full spatio-featural activity-space, we encoded DCNN activity into 1000 components using principle components analysis (left, orange). To isolate spatial activity, we averaged across the channel dimension in DCNN activity (center, blue). To isolate spatially invariant feature-based activity, we averaged across the spatial dimensions (right, green). **d**. Object category (binary prediction between all 28 combinations of the eight categories, chance = 50%) could be predicted from all types of DCNN activity, with components and feature-based activity showing a sharp rise in decodability across layers. This plot does not include any neural data, it only shows decodability using DCNN activity.

First, we defined an intermediate space based on unit activity in a DCNN trained for object categorization (VGG-16 pre-trained on ImageNet, **Supplemental Fig. 1**) [19]. For the pooling layers, we analyzed three features computed from DCNN activity: principal component scores, feature-based channel activity, and spatial activity (**Fig. 1c**). For the fully-connected layers, we analyzed two DCNN features: principal components and feature-based channel activity. Within each layer, each DCNN feature for each layer was mapped to object category using a series of 8-choose-2 linear support vector machine (SVM) classifiers to make binary predictions of object category. Importantly, the training of these SVMs did not incorporate any neural data. Object category could be predicted from each of the seven VGG-16 layers for all three DCNN features (**Fig. 1d**). For components and feature-based activity, and to a lesser extent for spatial activity, performance improved for progressively deeper layers relative to earlier layers.

Next, we mapped IT activity to DCNN activity using linear regression (20 category-matched 75% train, 25% test folds). The resultant IT to DCNN transformation matrices were multiplied by IT activity vectors from the test set to transform IT activity into the same space as DCNN component scores. The transformed IT activity was multiplied by the transpose of the PCA transformation matrix to reconstruct the full DCNN activity space for a given layer. As with true DCNN activity, feature-based and spatial reconstructions were computed for the five pooling layers, and the full 4096 reconstructed channel activations were analyzed for fully-connected layers. This procedure was repeated separately for each layer and cross-validation fold (**Supplemental Fig. 2**).

**Figure 2.**
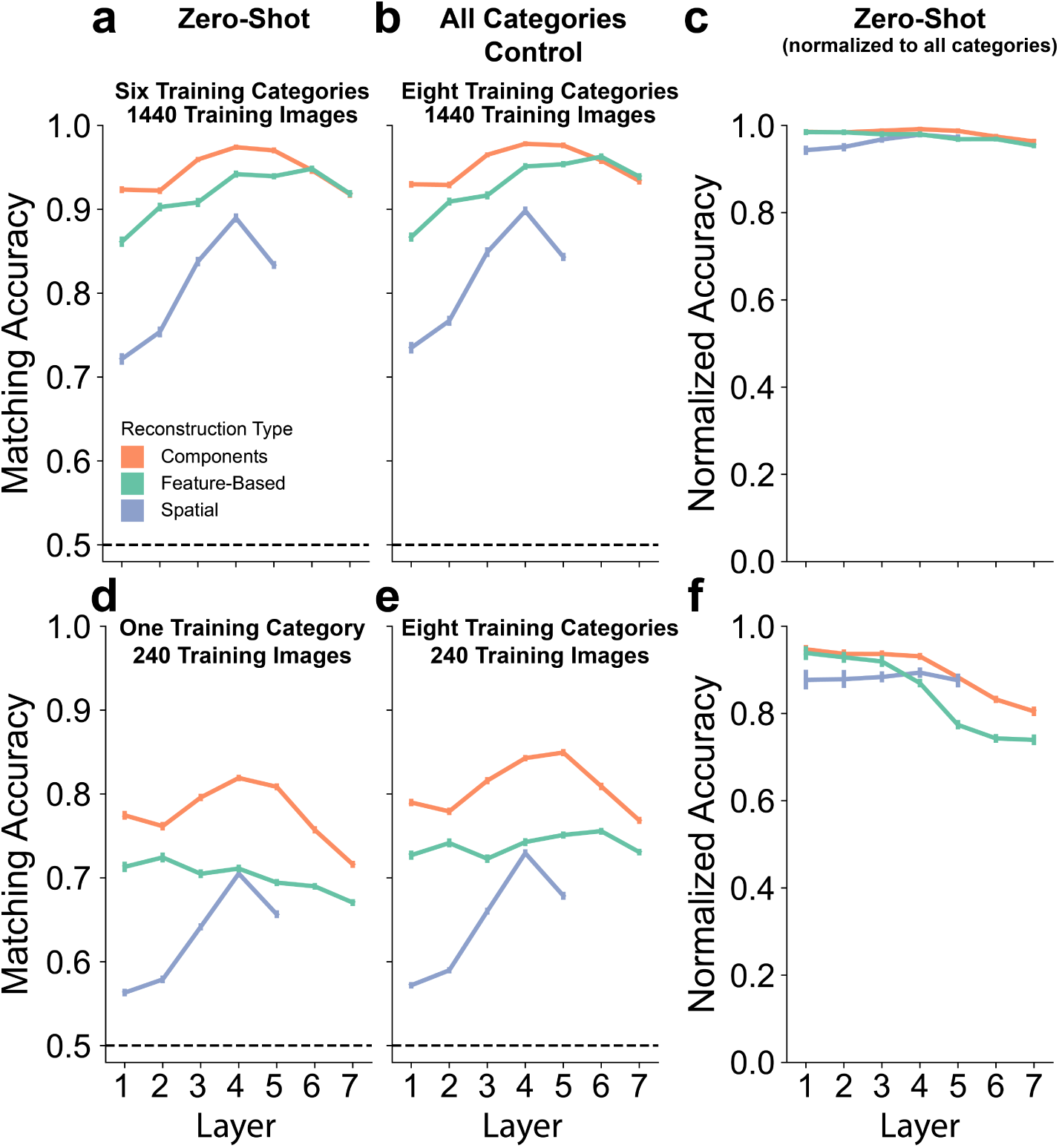
DCNN features reconstructed from IT activity match true DCNN features for the same images even when extrapolating across categories. **a**. High matching accuracies were achieved when neural activity from two test categories was held out during training (six training categories), indicating that IT to DCNN mappings indeed generalize across object category. **b**. As a control, a decoder was trained on neural responses from all eight categories, matching the overall number of training images to the number used for the zero-shot decoder. Strikingly similar results were obtained for the zero-shot and all categories control decoders. **c**. Zero-shot matching accuracies were normalized to calculate proportion of above-chance matching accuracy achieved by the zero-shot model relative to the all categories model. Normalized matching accuracies are close to ceiling for all layers and reconstruction types. **d**, **e**, **f**. Zero-shot, all categories control, and normalized results when only one category was used to train the zero-shot decoder. Again, the zero-shot decoder displayed highly similar results to the all categories control decoder. All accuracies for all decoders, DCNN feature-types, and layers are significant at P *<*0.001 (permutation testing).

Finally, to determine whether the mapping between IT and DCNN activity extrapolates to novel categories, IT activity from the test categories was held out during training of the mapping. If decoding accuracies when recognizing novel categories are high, this indicates that the IT to DCNN mapping is zero-shot, capturing generic visual information in IT to generalize to novel categories on which it was never trained.

To assess the overall amount of shared generic visual information between IT and DCNN activity, reconstructed DCNN features were matched to the true DCNN features from the same images, relative to the true DCNN features from every other image in the test set, for all possible pair-wise combinations. We measured the matching accuracy, which ranges from 0.5 (chance) to 1 (perfect reconstruction of features) (**Fig. 2**). We examined the extreme cases where the maximum number of available training data (six categories) and minimum number (one category) were used.

When the zero-shot decoder was trained on neural responses from six categories, matching accuracies for reconstructed DCNN components, feature-based activity, and spatial activity were all well above chance (all Ps <0.001, permutation testing), with components and feature-based activity exhibiting higher matching accuracies than spatial activity (**Fig. 2a**). As a control, we compared the zero-shot decoder’s performance to a decoder trained on responses from all categories, matching the overall number of training images to the number used for the zero-shot decoder. This all categories control decoder displayed strikingly similar results to the zero-shot decoder (**Fig. 2b**, all Ps <0.001, permutation testing). A normalized matching accuracy, the proportion of above-chance matching accuracies achieved by the zero-shot decoder relative to the all categories control decoder, was calculated as (matching accuracy_Zero-Shot_– chance) / (matching accuracy_AllCategories_ – chance). A normalized matching accuracy of 0 indicates zero-shot performance was at chance, a value of 1 indicates the zero-shot matching accuracy was equal to the matching accuracy for the model trained on all categories. Normalized matching accuracies were all close to ceiling (**Fig. 2c**), indicating strinkingly comparable accuracies between the zero-shot and all categories control decoders.

Even when the zero-shot decoder was trained on neural responses from just one category, matching accuracies remained well above chance (**Fig. 2d**, all Ps <0.001, permutation testing). Again, after computing normalized accuracies relative to an all categories control decoder matched for the number of training images (**Fig. 2e**), we find normalized accuracies close to ceiling (**Fig. 2f**), demonstrating strong generalization of the IT to DCNN mappings. Full matching results for all possible numbers of training categories can be seen in **Supplemental Fig. 3**.

**Figure 3.**
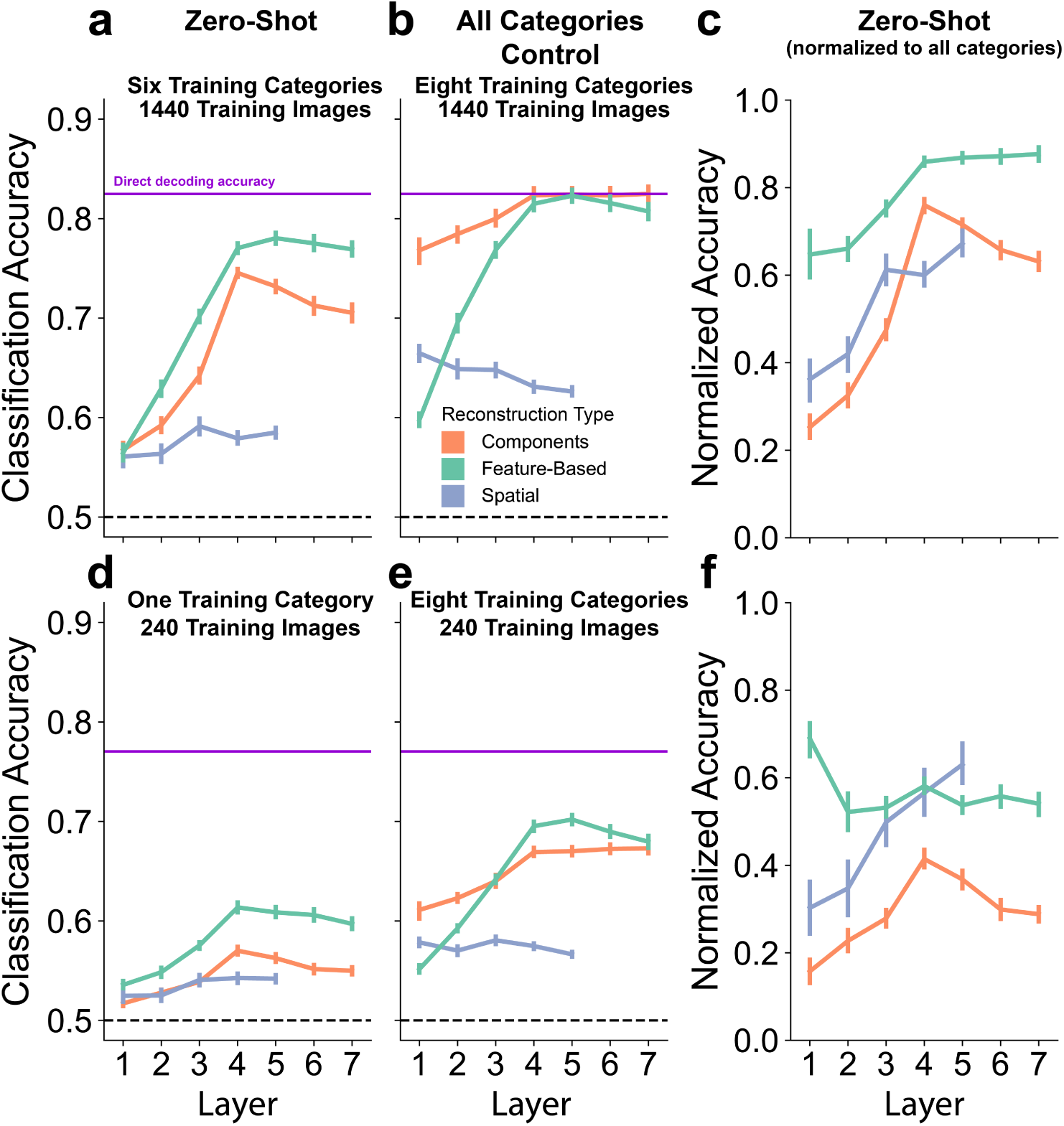
Object category can be decoded from IT-reconstructed DCNN features even when extrapolating to novel categories that were not used for training the IT to DCNN mapping. **a**. High classification accuracies were achieved when neural activity from two test categories was held out during training (six training categories), indicating that IT to DCNN mappings capture generic information about visual object category. Direct decoding accuracies (predicting category directly from IT responses using linear SVMs) are shown in purple. **b**. A control decoder, where the IT to DCNN mapping was learned using IT responses from all eight categories, showed a similar pattern of results, albeit with higher accuracies. **c**. Zero-shot classification accuracies were normalized to calculate proportion of above-chance classification accuracy achieved by the zero-shot model relative to the all categories model. Normalized classification accuracies were all greater than zero, and the best normalized accuracies for feature-based reconstructions achieved over 80% of the accuracies seen for the all categories control decoder. **d**, **e**, **f**. Zero-shot, all categories control, and normalized results when only one category was used to train the zero-shot decoder. Again, the zero-shot decoder displayed a similar pattern of results to the all categories control decoder but with lower accuracies. All accuracies for all decoders, DCNN feature-types, and layers are significant at P *<*0.001 (permutation testing).

Next, we assessed whether the information captured in the IT to DCNN mappings is discriminative of object categories. For this purpose, we used the same 8-choose-2 SVM classifiers, trained on DCNN activity (**Fig. 1d**), *without retraining or fine-tuning on neural data*, to generate object category predictions from IT-reconstructed DCNN features (**Fig. 3**). In other words, the model presented with IT responses from novel categories and the task is decode which category was presented to the monkey, even though the mapping to reconstruct DCNN activations from IT was never exposed to neural responses from that particular category.

When the zero-shot decoder was trained on six categories, we see significant prediction for all reconstructed DCNN features and layers (**Fig. 3a**, all Ps <0.001, permutation testing). Feature-based reconstructions produced the best zero-shot predictions and were well above chance. The feature-based reconstructions show an increase in zero-shot prediction accuracy over the first few layers before flat-lining, indicating an expected increase in shared information about object category between IT and the DCNN across layers. Again, we compare the zero-shot performance to a control decoder where the IT to DCNN mapping was learned on responses from all categories (**Fig. 3b**, all Ps <0.001, permutation testing)) by calculating normalized classification accuracies. These normalized classification accuracies were all well above 0, with the feature-based reconstructions from the later layers achieving a proportion over 0.8 of the accuracies seen for the all categories decoder (**Fig. 3c**). These results demonstrate that information captured about the neural code for object representation when learning the mapping from IT to DCNN activity extrapolates to untrained novel object categories.

When a single category was used to train the zero-shot decoder, accuracies were still all significantly above chance (**Fig. 3d**, all Ps <0.001, permutation testing). After normalizing these accuracies to an all categories control model matched for training set size (**Fig. 3e**, all Ps <0.001, permutation testing), normalized accuracies were all greater than zero (**Fig. 3f**). Despite only ever being exposed to neural responses from a single object category, this zero-shot decoder was still able to make pair-wise category judgments for neural responses from seven held-out novel categories, displaying extreme generalization not previously reported for any neural decoding model from any imaging modality. Full classification results for all possible number of training categories can be seen in **Supplemental Fig. 4**.

Overall, mappings from IT to DCNN activity generalized across object category and novel object categories could be predicted without the model having prior exposure to responses from those categories, providing evidence that DCNNs capture generic visual information in *rhesus macaque* IT, as opposed to information that is restricted to the categories used for fitting. Understanding the neural code for objects requires not only *interpolation* to novel test items similar to those in the training set (as is standard practice), but also *extrapolation* to completely novel shapes that are clearly distinct from those in the training set. Such zero-shot generalization demonstrates that the model has captured the inherent structure of information encoded in neural activity and the relationship between the encoded features and object category. The extreme case of successful generalization from just a single training category (**Figs. 2d,f and 3d,f**) suggests a robust relationship between IT and DCNN representations.

In studies linking DCNN features to brain activity, the DCNN units are usually treated equally without regard to the native dimensions in DCNN representations. Here, we separate feature-based and spatial activity, as well as principal component scores summarizing the full spatio-featural activity space, to provide greater clarity regarding which aspects of DCNN representations are explaining the neural variance that carries information about object category. We find that feature-based DCNN channel activity reconstructed from IT activity carries the most generic visual information.

This work demonstrates the promise of zero-shot neural decoders from electrophysiological recordings, which has broad applications. Theoretically, zero-shot decoders are superior to standard decoders because they necessitate a computational hypothesis for the neural code underlying the targeted process and support maximal generalization. On the engineering side, they could enable advances such as decoders from chronic neural recordings that can be flexibly updated without new training data. Rather than directly mapping neural responses to every desired output, neural activity would be mapped to a convenient intermediate space that captures the relevant variance for many different outputs. Predicting novel information from the decoder would simply necessitate training a new computational model linking the intermediate features to the new outputs, rather than collecting new neural recordings to learn a direct mapping. As chronic neural recordings become commonplace, building such flexible, generalizable neural decoding systems will become ever more important.

## Methods

### Dataset

Details about the experimental setup, recording procedure, and pre-processing can be found in [18]. Briefly, two awake *rhesus macaque* monkeys were passively shown a rapid-serial-visual-presentation stream of 2560 grayscale images, each presented 50 times (28 minimum repetitions) for 100 ms, depicting computer-generated objects from eight categories (Airplanes, Animals, Boats, Cars, Chairs, Faces, Fruits Tables) superimposed on arbitrary natural scene backgrounds. Within each of the eight categories, there were eight unique objects (40 images per object). The full stimulus set in [18] had three conditions of objects: low-variation (same size, position, orientation across all background), medium-variation (some variation in size, position, orientation across backgrounds), and high-variation (high variation in size, position, orientation across backgrounds). Here, only images and IT responses from the high-variation condition were used (in which behavioral recognition for a monkey or machine would be most difficult). Neural recordings were acquired from 168 visually-selective IT units using multi-electrode arrays. Firing rates were calculated from 70 to 170 ms post stimulus onset and averaged across repetitions.

### Cross-Validation Folds

For all analyses, data were split into category-matched training and testing folds. Data were split into folds according to object, so even within-category the specific objects used in the various training and testing phases were independent. DCNN activity from 75% of the images (six objects per category) was used to map DCNN activity to category labels, define the PCA transform on DCNN activity, and learn the mapping between IT and DCNN activity (Fig. 1). Neural and DCNN activity from 25% of the images (two objects per category) was used to test the models. Twenty unique train-test splits were used for all analyses.

### DCNN Architecture

We used VGG-16 trained for 1000-way object categorization on the ImageNet dataset [19]. VGG-16 is a hierarchical DCNN with 21 convolutional, max-pooling, and fully-connected layers (**Fig 1a**. middle column). Our analyses focused on the five pooling layers (pool 1 = 802,816 units, pool2 = 401,408 units, pool3 = 200,704 units, pool4 = 100,352 units, pool5 = 25,088 units, fc6 = 4096 units, fc7 = 4096 units), which were selected to sample DCNN activity from across the entire hierarchy. The full unit activity space for each layer was reduced to 1000 principle component scores using PCA (75% train, 20% test cross-validation splits). To isolate feature-based channel activity in the pooling layers, we averaged units across the spatial dimensions (pool 1 = 64 channels, pool2 = 128 channels, pool3 = 256 channels, pool4 = 512 channels, pool5 = 512 channels). To isolate spatial activity in the pooling layers, we averaged unit activity across channels (pool1 = [112, 112], pool2 = [56, 56], pool3 = [28, 28], pool4 = [14, 14], pool5 = [7, 7]. The fully-connected layers are organized along a single channel dimension, so all 4096 units were included as feature-based activity.

### DCNN Readout SVMs

The relationship between DCNN activity and object category labels was learned using linear support-vector-machines (SVMs). DCNN activity from 75% of the images (six objects/category) were used to train the SVMs, and DCNN activity from 25% of the images (two different objects/category) were used to test the SVMs. Prior to training, each unit was normalized to have a mean of zero and a standard deviation of one across images. The same scaling learned on the training set was applied to the test set. Twenty-eight 8-choose-2 binary SVMs were trained, one for every potential pair amongst the eight object categories in the dataset. Binary classification was selected so the two test categories could easily be held-out in the zero-shot condition. Hyper-parameters for each binary classifier were optimized to maximize classification accuracy using three-fold cross-validation within the training set. The same DCNN readout SVMs were used to predict object category from DCNN activity for all analyses. To emphasize, the DCNN readout SVMs were trained independent from any electrophysiological recordings and were never exposed to IT activity until the test phases.

### Decoding Methodology

We used partial least squares regression (PLSR) with 25 components (as in [13]) to learn the mapping between IT activity and DCNN activity. IT and DCNN activity from 75% of the images (six objects/category) were selected as the training set to learn the IT to DCNN transformation. When IT and DCNN activity from all eight categories were used to learn the transformation, the same transformation was applied to test IT activity from all categories. For zero-shot decoding, one to six categories were used to learn the transformation, and the transformation was applied to the held-out test categories. To assess how well the IT to DCNN transformation extrapolates to novel categories in the most extreme conditions, we used all possible numbers of categories (one to six, step-size one) to learn the IT to DCNN transformation. In all of the above versions, the IT activity transformed into DCNN activity was passed into the DCNN readout SVMs to generate the final object category predictions.

### Matching Analysis

To assess the accuracy of the reconstructions and obtain a measure of the overall amount of shared generic information between IT and DCNN activity, we matched DCNN features reconstructed from IT to true DCNN features from the same image. In a pairwise fashion, the reconstructed DCNN features were correlated (Pearson) with the true DCNN features for the same image and the true DCNN features from another image in the test set. If the within-image correlation is greater than the between-images correlation, that comparison was scored as a hit. For a given target image, this comparison was made for every other image in the test set, and the same procedure was applied using each test image as the target image. We averaged across all comparisons to get a matching accuracy (50% chance). Significance was determined using permutation testing. Reconstructed DCNN features were permuted 1000 times relative to their image labels and the full analysis was run to derive a null distribution of matching accuracies. P was defined as the proportion of matching accuracies from this null distribution that are greater than the true matching accuracy. The matching analysis was run for each DCNN feature-type and layer.

### Classification Analysis

To assess whether DCNN features reconstructed from IT activity contain information about object category, the reconstructions were fed into the 8-choose-2 linear SVMs trained on true DCNN activity. The SVMs were not modified or fine-tuned on any neural activity. Significance was derived using permutation testing, again permuting the reconstructed DCNN features relative to image labels to derive empirical null distributions of classification accuracies. The classification analysis was run for each DCNN feature-type and layer.

### Normalizing Zero-Shot Accuracies

To better assess zero-shot accuracies and compare across conditions with different numbers of training categories, we calculated normalized zero-shot accuracies. The scores account for the proportion of above-chance accuracy present in a zero-shot decoder relative to a control decoder trained on neural responses for all categories. In all comparisons, the number of training images for the all categories control decoder was matched to the number of training images for the zero-shot decoder.

## Acknowledgments

This work was supported by the Center for Brains, Minds and Machines, funded by NSF Science and Technology Centers Award CCF-1231216, aand NIH Grant R01EY026025. M.M.C is funded by NIMH 108591. We thank Jim DiCarlo for sharing the IT recording data used in this experiment. We thank Tyler Bonnen, Kasper Vinken, Yaoda Xu, and Daeyeol Lee for helpful comments on this work.

## Author Contributions

T.P.O’C. conceived of the study. T.P.O’C. and G.K. designed the study. T.P.O’C. performed the analyses. T.P.O’C. and G.K. wrote the manuscript with contributions from M.M.C.

---

**Figure S1.**
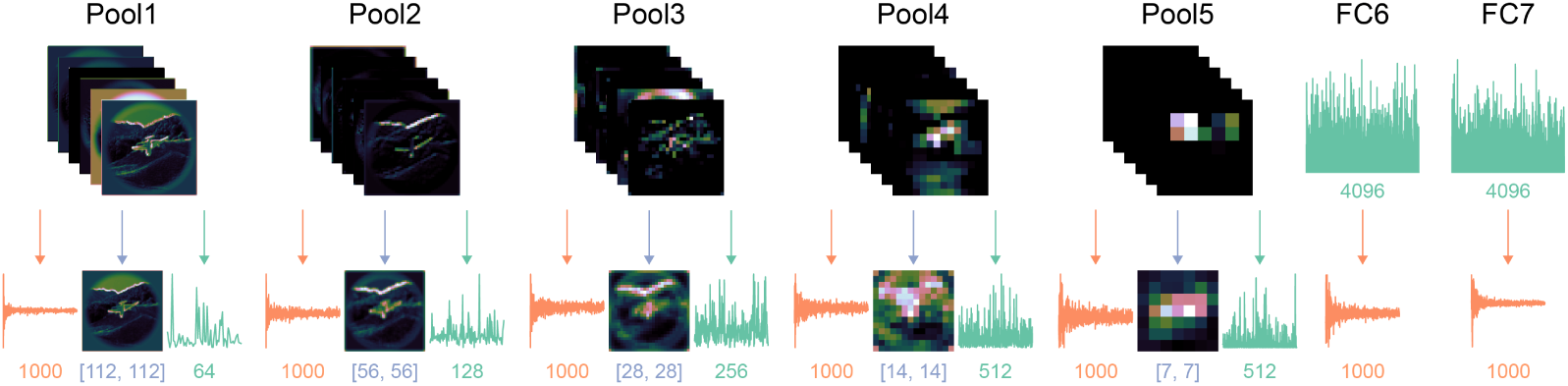
Features extracted from each layer of VGG-16. Components (orange) were defined using principle components analysis and the number of components (1000) was matched across layers. Spatial activity (blue) was defined by averaging across the channel dimension in the native unit-activity space. Feature-based activity (green) was defined by averaging unit activity across the spatial dimensions. The final dimensionality of each feature-type is shown.

**Figure S2.**
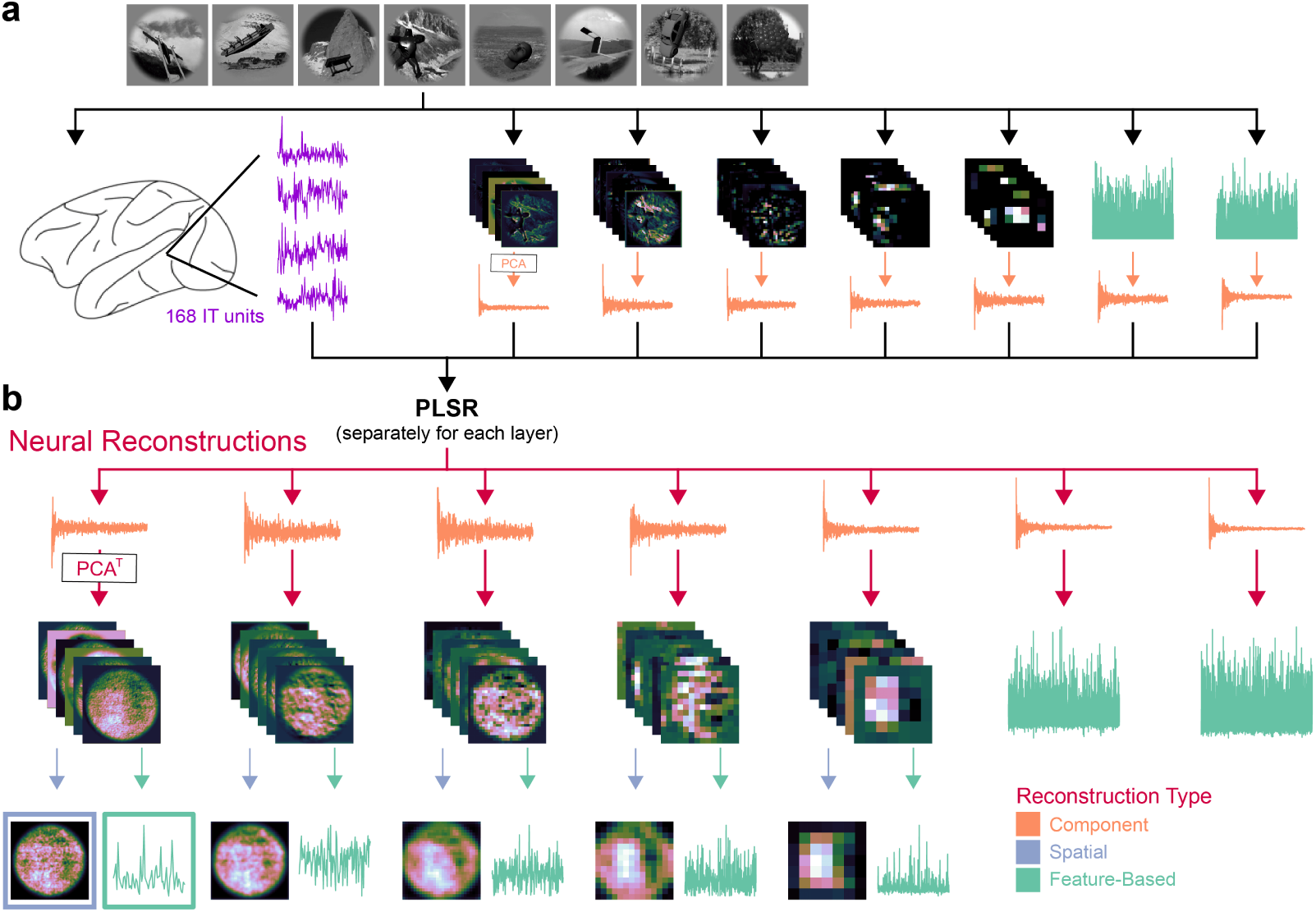
a. Two rhesus macaque monkeys viewed images from 8 object categories while IT responses were recorded using multi-electrode arrays. DCNN activity for each image was computed using VGG-16, and the full unit activity for each layer was encoded into 1000 components using PCA. **b**. Using partial least squares regression (PLSR), linear mappings were learned from IT response patterns to DCNN components. These mappings were applied to held-out data (twenty 75% train, 25% test splits) to decode DCNN components from IT activity. The decoded components were multiplied by the transpose of the PCA transformation to reconstruct the full space of DCNN activity for each layer. In the five pooling layers, full reconstructions were averaged across channels (blue arrows) to calculate spatial reconstructions and across spatial dimensions (green arrows) to calculate spatially invariant feature-based reconstructions.

**Figure S3.**
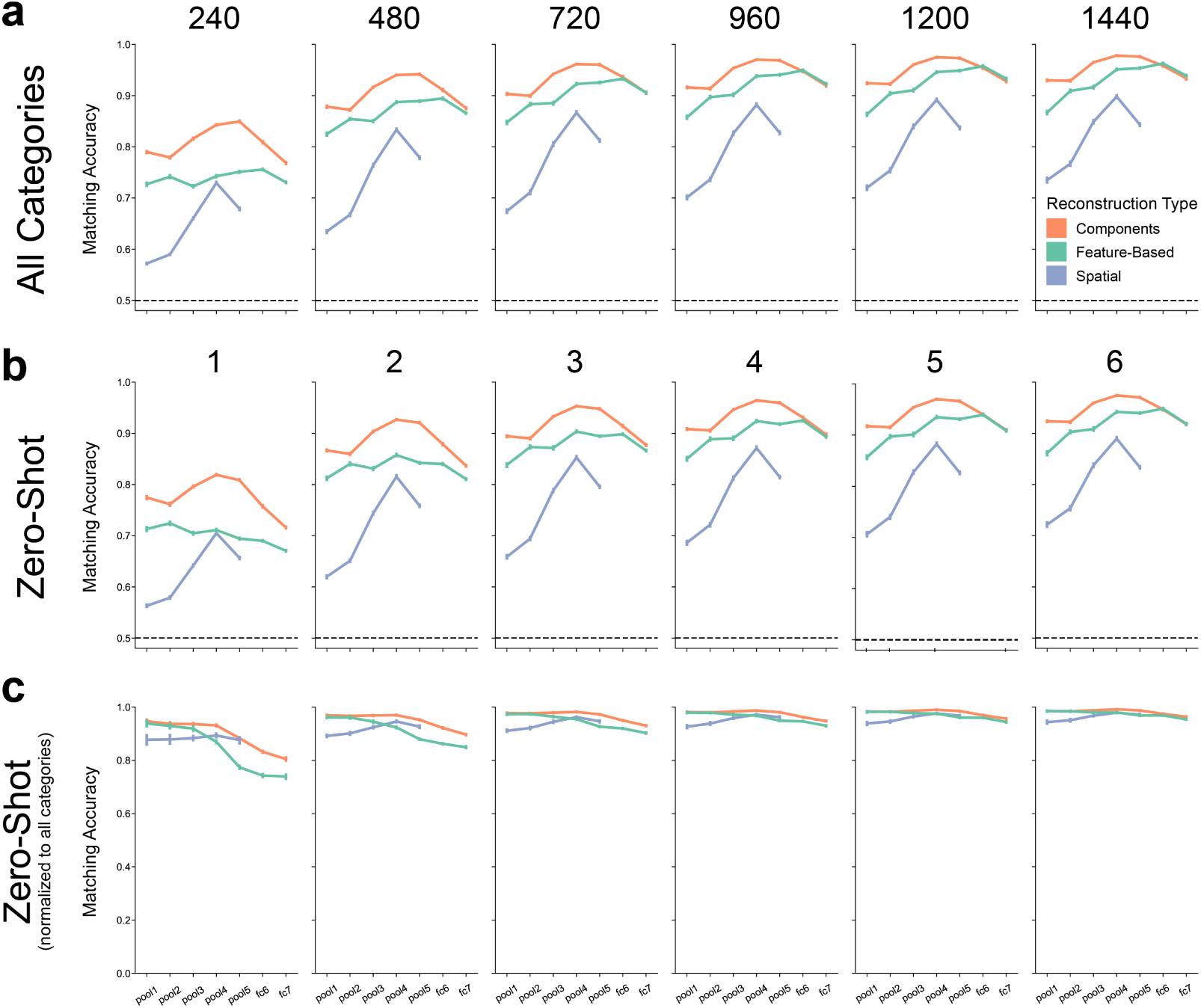
(expanding on Fig. 2) Matching results for all possible number of training categories. **a**. Model-based matching accuracy when training on neural activity from all categories (similar to **Fig. 2**). The number of training images, shown along the top of each column, are matched to the number of training images for each zero-shot condition below. The subplot for 240 training images corresponds to **Fig. 2e** and the subplot for 1440 training images corresponds to **Fig. 2b. b**. Zero-shot matching accuracy for all possible number of training categories. The number of training categories are shown at the top of each column. The subplot for one training category corresponds to **Fig. 2d** and the subplot for six training categories corresponds to **Fig. 2a. c**. Zero-shot matching accuracy normalized by the all categories matching accuracy. The subplot for one training category corresponds to **Fig. 2f** and the subplot for six training categories corresponds to **Fig. 2c**.

**Figure S4.**
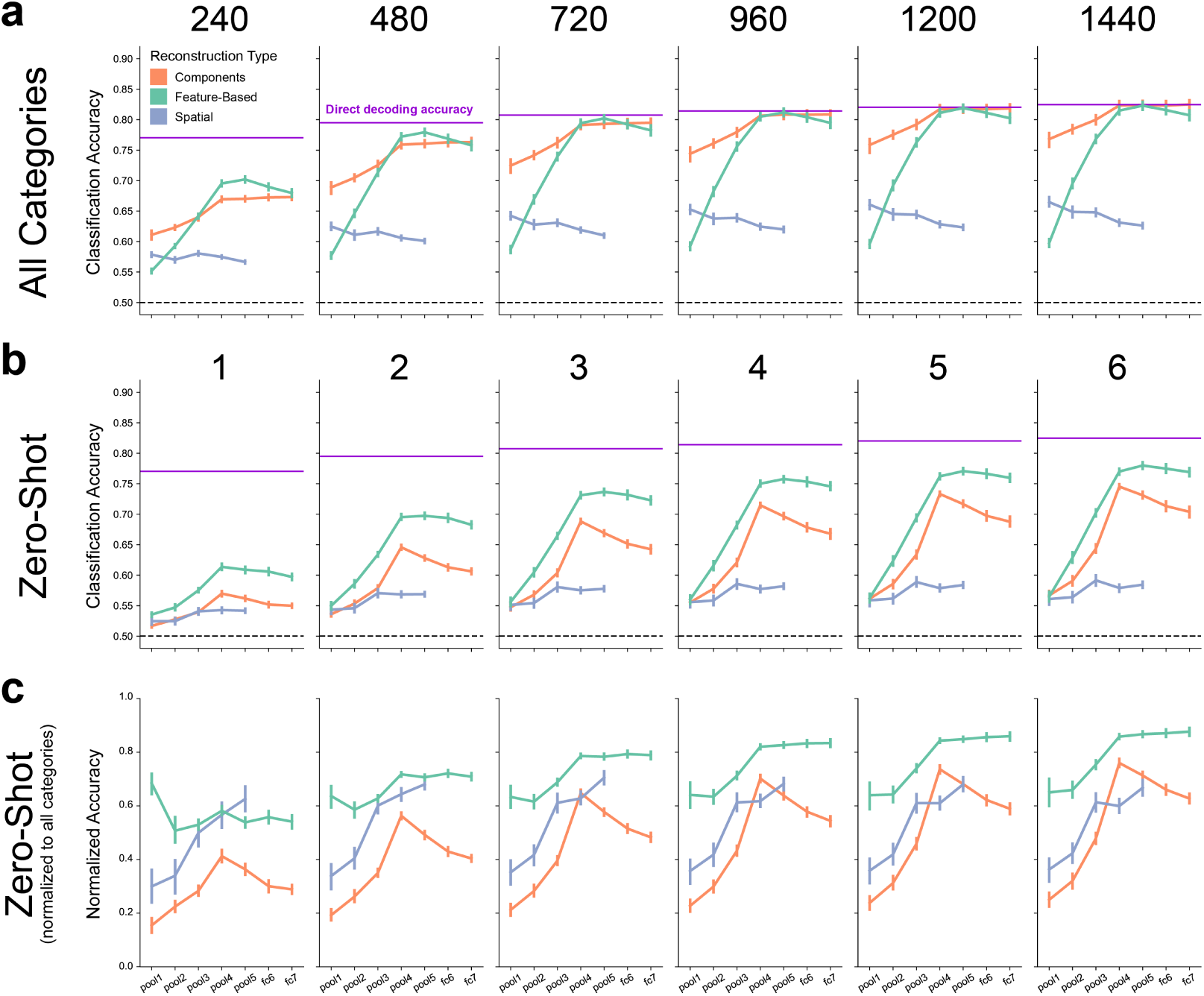
(expanding on Fig. 3) Classification results for all possible number of training categories. **a**. Model-based classification accuracy training on neural activity from all categories (similar to **Fig. 3**). The number of training images, shown along the top of each column, were matched to the number of training images for each zero-shot condition below. The subplot for 240 training images corresponds to **Fig. 3e** and the subplot for 1440 training images corresponds to **Fig. 3b. b**. Zero-shot classification accuracy for all possible number of training categories. The number of training categories are shown at the top of each column. The subplot for one training category corresponds to **Fig. 3d** and the subplot for six training categories corresponds to **Fig. 3a. c**. Zero-shot classification accuracy normalized by the all categories classification accuracy. The subplot for one training category corresponds to **Fig. 3f** and the subplot for six training categories corresponds to **Fig. 3c**.

